# Species-specific roles for the MAFA and MAFB transcription factors in regulating islet β cell identity

**DOI:** 10.1101/2022.10.20.512880

**Authors:** Jeeyeon Cha, Xin Tong, Emily Walker, Tehila Dahan, Veronica Cochrane, Sudipta Ashe, Ronan Russell, Anna Osipovich, Alex Mawla, Min Guo, Jin-hua Liu, Mark Huising, Mark Magnuson, Matthias Hebrok, Yuval Dor, Roland Stein

## Abstract

Type 2 diabetes (T2D) is associated with compromised identity of insulin-producing pancreatic islet beta (β) cells, characterized by inappropriate production of other islet cell-enriched hormones. Here we examined how hormone misexpression was influenced by the MAFA and MAFB transcription factors, closely related proteins that maintain islet cell function. Mice specifically lacking MafA in β cells demonstrated broad, population-wide changes in hormone gene expression with an overall gene signature closely resembling islet gastrin (Gast)-positive cells generated under conditions of chronic hyperglycemia and obesity. A human β cell line deficient in MAFB, but not one lacking MAFA, also produced a gastrin (GAST)-positive gene expression pattern. In addition, GAST was detected in human T2D β cells with low levels of MAFB. Moreover, evidence is provided that human MAFB can directly repress *GAST* gene transcription. These results support a novel, species-specific role for MafA and MAFB in maintaining adult mouse and human β cell identity, respectively, by repressing expression of Gast/GAST and other non-β cell hormones.

## INTRODUCTION

The pancreatic islet is comprised of endocrine cells expressing distinct peptide hormones, such as insulin from β cells, glucagon from α cells, and somatostatin from δ cells, which play critical roles in maintaining euglycemia. Experiments conducted principally in mouse models have demonstrated that combinatorial expression of islet-enriched transcription factors (TFs), such as Pdx1, Nkx6.1, Arx, Hhex and MafA are essential in the developmental and/or adult function of islet cell types (1, 2). For example, studies performed on *Maf<u>a</u>* mutant mice (global *MafA*^−/-^, pancreas-specific *MafA*^*Δpanc*^, and β cell-specific *MafA*^*Δβ*^) established a key role for this TF in activating programs promoting β cell maturation and glucose-stimulated insulin secretion (GSIS) (1, 3). In contrast, the related MafB protein is present throughout the lifespan of the α cell and has limited importance to adult islet β cells: this TF is only produced in murine β cells embryonically, except when transiently expressed during pregnancy to facilitate β cell expansion (4). Moreover, MafB cannot rescue MafA function in MafA-deficient β cells (4), likely due to unique coregulator(s) interactions within their distinct C-terminal region sequences (5).

While expression of most islet-enriched TFs is similar between rodents and humans, the MAFA and MAFB proteins have very different temporal profiles in humans. MAFB is expressed throughout the lifespan of both human α and β cells, which contrasts from the postnatal rodent MafA^+^;MafB^−^ β cell population (1). In addition, MAFA is not readily detectable in human islet β cells until ∼9 years of age (4), while in rodents, this TF is expressed at the onset of insulin^+^ cell formation during embryogenesis (6). In contrast to rodents, human MAFB appears to be essential for β cell development, as supported by the inability of human embryonic stem cells (hESCs) lacking MAFB to produce insulin (INS^+^) cells despite being subject to a well-established β cell differentiation protocol (7). Strikingly, these mutant cells still produce many islet-enriched TFs associated with *INS* transcription (e.g., NKX6.1 and PDX1). In addition, a subpopulation of MAFB^KO^ β-like cells express non-β cell hormones, such as somatostatin (SST), pancreatic polypeptide (PPY), and GAST, with the latter principally expressed in adult gastric (G) cells and only transiently expressed in the pancreas during islet formation embryonically (8).

Since synthesis of non-β hormones appears to be negatively regulated by human MAFB (7) and is associated with the loss of β cell identity and activity postnatally in T2D (9-13), we examined here how MAFA and/or MAFB contributed to this process in adult mice and human islets. Non-β cell hormone-producing cells were not only inappropriately generated in mouse *MafA*^*Δβ*^ islets but were also detected under pathophysiologic hyperglycemic and obesity conditions that suppress expression of this TF. Notably, the gene expression profile of mouse islet gastrin (Gast^+^) cells produced in hyperglycemic models was shared with the broader *MafA*^*Δβ*^ β cell population, even though only a small fraction of *MafA*^*Δβ*^ islet cells were Gast^+^. SST, PPY, and GAST were also inappropriately expressed upon knockdown of MAFB, but not MAFA, in the human EndoC-βH2 β cell line. Furthermore, both SST^+^ and GAST^+^ cells were only observed in the MAFB^Low^ islet β cells in human T2D islets. Because MAFA and MAFB protein levels are more sensitive to metabolic and oxidative stress than other islet-enriched TFs (14), we discuss the possibility that their loss under such conditions contributes to changes in β cell identity and inactivity associated with T2D progression.

## RESULTS

### Non-β cell hormone expression is induced in mouse *MafA*^*Δβ*^ islets

Our earlier results had shown that differentiation of the male human embryonic MEL-1 stem cell line to β-like cells requires MAFB, with MAFB-deficient β-like cells having unanticipated features including loss of INS production and generation of cells producing several non-β cell hormones (7). To determine if MafA had similar regulatory properties in postnatal mouse islet β cells, a β cell-specific deletion mutant was generated by crossing *MafA*^*fl/fl*^ mice (3) with *Ins1-Cre* mice (15), which effectively removed this TF from male and female islet β cells (termed *MafA*^*Δβ*^) (**Supplemental Figure 1A-B**). Notably, this *Ins1-Cre* line allows deletion only in the pancreatic β cell lineage, unlike other *Insulin* enhancer/promoter-driven Cre recombinase mouse lines that misexpress the transgene to drive recombination in the brain (16). As reported earlier for *Mafa*-deficient mouse lines, male and female *MafA*^*Δβ*^ mice similarly manifested impaired glucose tolerance without overt hyperglycemia (3, 4, 6, 17, 18).

**Figure 1:**
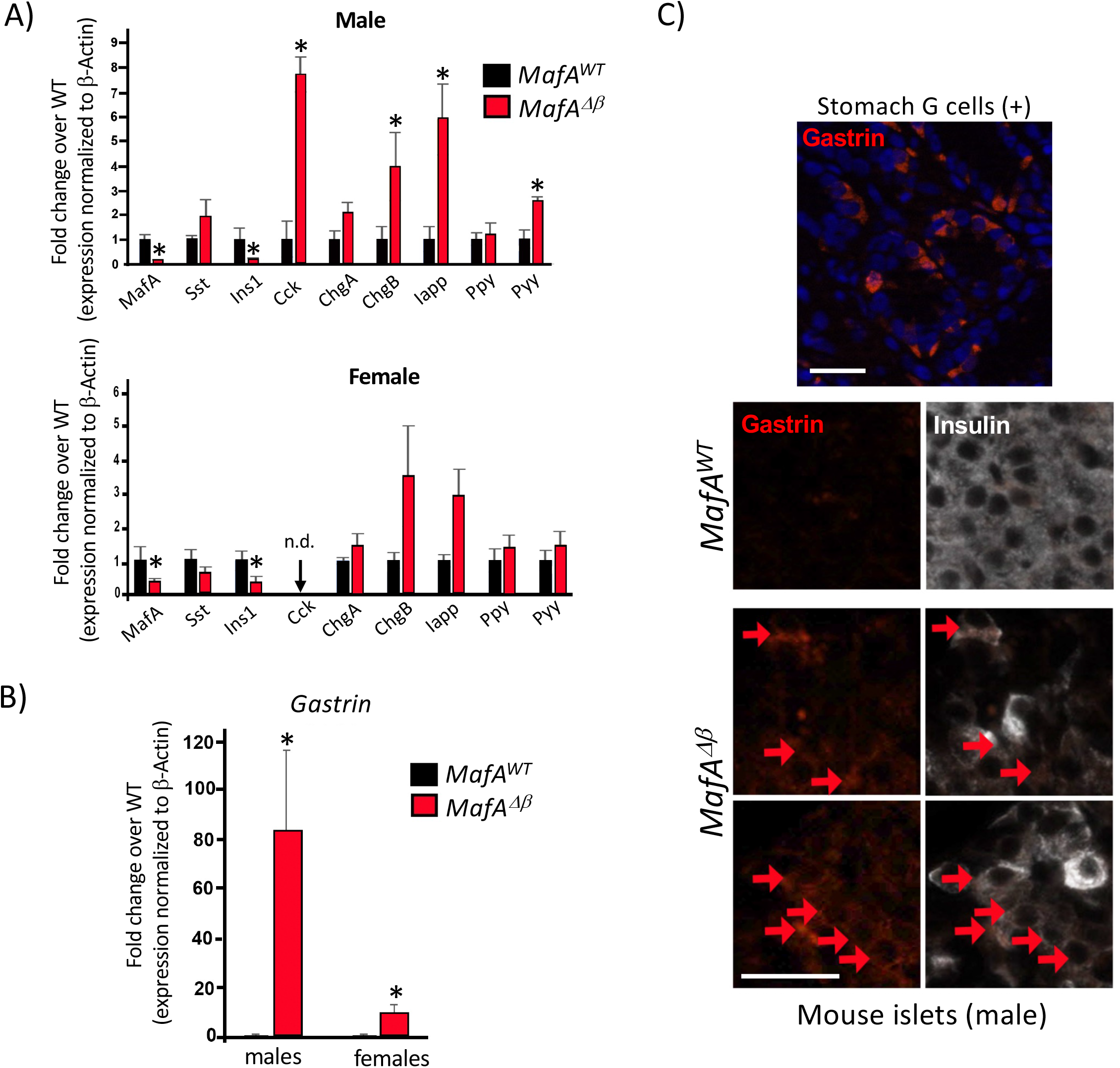
Non-β cell hormone expression is increased in *MafA*^*Δβ*^ islets. (A, B) Analysis of *MafA*, insulin 1 (Ins1) and non-β cell hormone mRNA levels in *MafA*^*WT*^ and *MafA*^*Δβ*^ islets from 3-month-old adult mice. (B) *Gast* mRNA was induced more in male, than female, *MafA*^*Δβ*^ islets. Mean ± SEM. n=3-4 animals/group. *p<0.05; n.d., not detectable. (C) Gast (red) and Insulin (white) immunostaining in male *MafA*^*WT*^ and *MafA*^*Δβ*^ islets. The subpopulation of co-producing cells is denoted by red arrows. Stomach G cells served as a positive control for Gast staining. n=3-5 animals/group. Bar, 50µm.

Analysis of endocrine cell hormone mRNA expression in FACS-purified β cells from adult *MafA*^*Δβ*^ islets revealed that *Insulin (Ins)* levels were reduced, while expression of non-β cell hormones such as *cholecystokinin* (*Cck*), *chromogranin B* (*Chgb*), *Gast* and *Pyy* were elevated (**Figure 1A-B**). Interestingly, a profound sexual bias was observed for *Cck* (**Figure 1A**) and *Gast* (**Figure 1B**). Gast protein was also detectable in male and female *MafA*^*Δβ*^ islets but not *MafA*^*WT*^ (**Figure 1C**). However, gene markers associated with β cell dedifferentiation, such as *Ngn3, FoxO1*, and *Aldh1a3* (19, 20), were not detectable in *MafA*^*Δβ*^ or control β cells in either sex (data not shown). Collectively, these results suggest that MafA can repress production of hormones that are normally only transiently expressed during pancreatic β cell development and predominately made elsewhere postnatally, such as *Gast* (duodenum, stomach) (8) and *Cck* (small intestine) (21).

### The gene signature of the Gast^+^ cells produced in male insulin-resistant mice is found within the *MafA*^*Δβ*^ β cell population

Administration of the insulin receptor antagonist S961 induced hyperglycemia in male mice (**Figure 2A**) and diminished islet β cell MafA protein and mRNA levels (**Figure 2B and data not shown**). S961 treatment also stimulated the production of a rare Gast^+^ cell population, which was characterized by single cell RNA sequencing (**Figure 2C**). Twenty genes were significantly enriched in Gast^+^ cells compared to Gast^−^ cells, including the *Gcg, Sst, Cck, Chga, Chgb, Pyy* and *Iapp* endocrine markers (**Supplemental Table 1**). While female islets from S961-treated mice were not analyzed here, the purified β cell population from both male and female islets were analyzed in *ob/ob* mice, a T2D model that manifests obesity, hyperglycemia, elevated plasma insulin (22) and loss of β cell MafA (23). The male, but not female, *ob/ob* β cell population expressed many of the genes present in male S961-induced Gast^+^ cells (**Supplemental Figure 2A**). (Huang, Mawla et al, *in preparation*).

**Figure 2:**
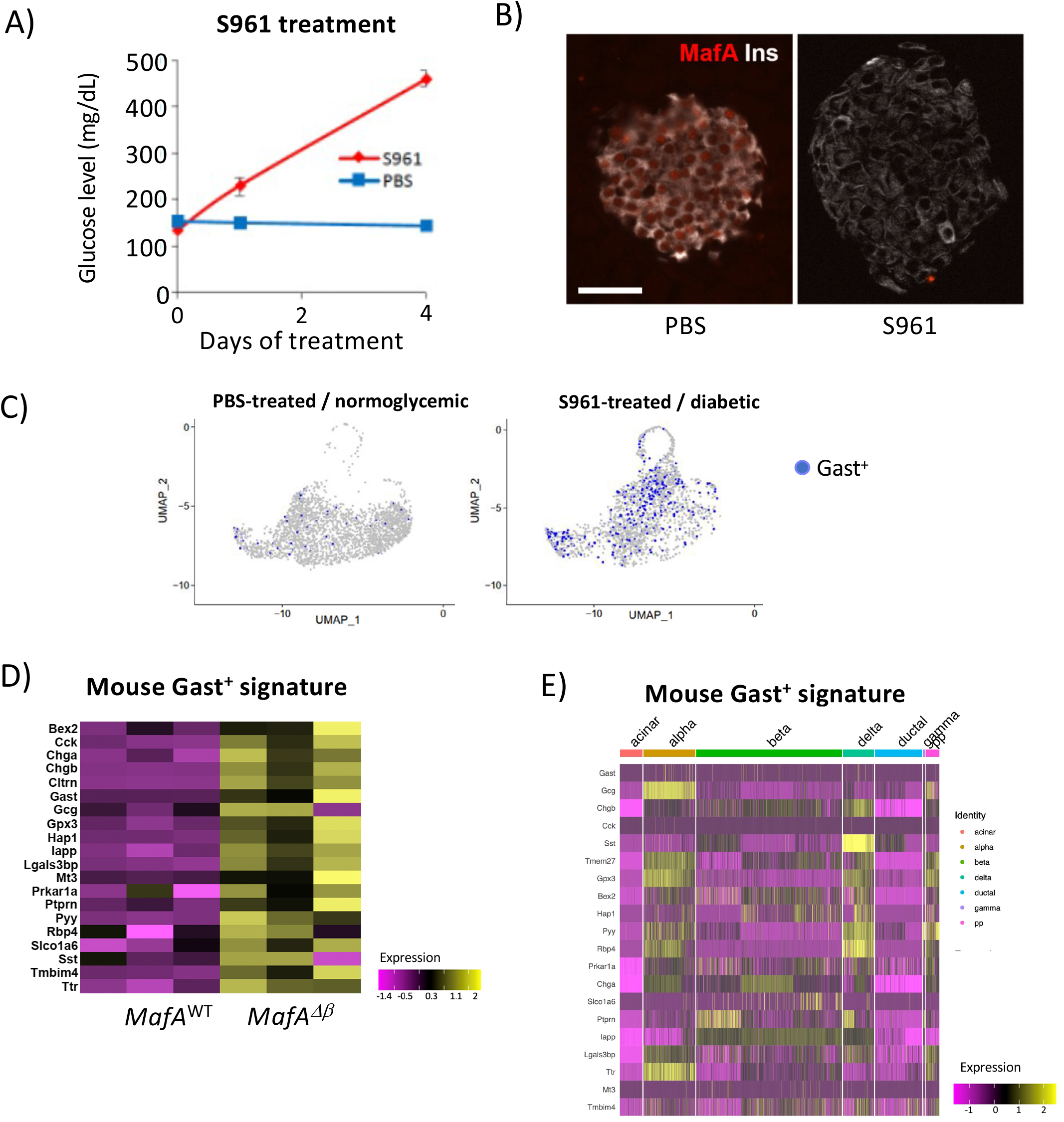
Islet Gast^+^ cells produced in insulin-resistant mice share molecular signatures with the broader *MafA*^*Δβ*^ islet population. (A) Administration of the S961 insulin receptor antagonist elevated random blood glucose levels in relation to PBS-treated control mice. (B) MafA (red) protein levels were markedly reduced in S961-treated mice after 7 days of treatment. Insulin (Ins, white). Bar, 50µm. (C) UMAP analysis of the single-cell RNA-seq data in male S961-treated islets shows increased number of Gast^+^ cells (blue dots) after 4 days of treatment. (D) The 20 up-regulated genes found in S961 Gast^+^ cells by single cell sequencing (**Supplemental Table 1**) were also elevated in male *MafA*^*Δβ*^ islets. *MafA*^*Δβ*^ β cell RNA-seq data was used in this analysis. n=3 animals/group. (E) There was limited overlap between the up-regulated genes found in S961 Gast^+^ cells and other exocrine and islet cell types.

In addition to *ob/ob* islet β cells, the genes associated with S961-treated mouse islet Gast^+^ cells were also enriched upon RNA-sequencing analysis of FACS-sorted male *MafA*^*Δβ*^ islet β cells (**Figure 2D**). Importantly, while the S961-induced Gast^+^ gene signature included several genes enriched in other islet cell types (i.e., *Gcg, Sst, Cck, Chga, Chgb, Pyy* and *Iapp*), there was little likeness at the gene expression level with naturally resident islet α, δ, and PP cells (**Figure 2E, Supplemental Table 1**) (24). While Gast itself is not broadly produced in male *MafA*^*Δβ*^ islet β cell population (**Figure 1C**), our results indicate that these cells express genes associated with the Gast^+^ cells generated in hyperglycemic and obese mouse models. Unfortunately, mouse single cell transcriptome data is currently unavailable for the Gast^+^ cells produced normally in the developing pancreas or more predominant stomach and duodenum G cells for comparison to the pathological gene signatures identified here in S961 and *ob/ob* islets. Of note, and as expected from analysis of hormone production in *MafA*^*Δβ*^ islet β cells (**Figure 1**), female *MafA*^*Δβ*^ β cells had less similarity at the gene expression level to these Gast^+^ cells than their male counterpart (**Figure 2D, Supplemental Figure 2B**).

### MAFB, and not MAFA, prevents *GAST, SST*, and *PPY* gene expression in human β cells

Adult human islet β cells express both MAFA and MAFB, while only MafA is produced in adult mouse β cells (5). Moreover, MAFA is not expressed until after the juvenile period in humans (4). To provide insight into the role of these proteins in controlling non-β cell hormone expression in human β cells, MAFA and MAFB knockdown experiments were performed in the MAFA/B- and MAFB only-expressing EndoC-βH1 (25) and EndoC-βH2 (26) β cell lines, respectively.

As predicted from the analysis of hESC-derived β-like cells lacking MAFB (MAFB^KO^) (7), lentivirus-mediated knockdown of MAFB protein (MAFB^KD^) in EndoC-βH2 β cells (**Figure 3A, left**) elevated *GAST, SST*, and *PPY* (increased trend) mRNA levels and decreased *INS* (**Figure 3A, right, gray bars**). An analogous regulatory pattern was observed upon knockdown of MAFB in EndoC-βH1 cells (**Figure 3B, right, grey bars**). Notably, immunostaining results showed that only a small fraction of MAFB^KD^ EndoC-βH2 cells express GAST (∼2%) or SST (∼2%) (**Figure 3C**). GAST and SST (GAST^+^SST^+^) co-positive cells were also observed in this setting (**Figure 3C**), similar to T2D islets (27). As expected, no shControl-treated EndoC-βH2 cells produced the GAST or SST protein (**Figure 3C**).

**Figure 3:**
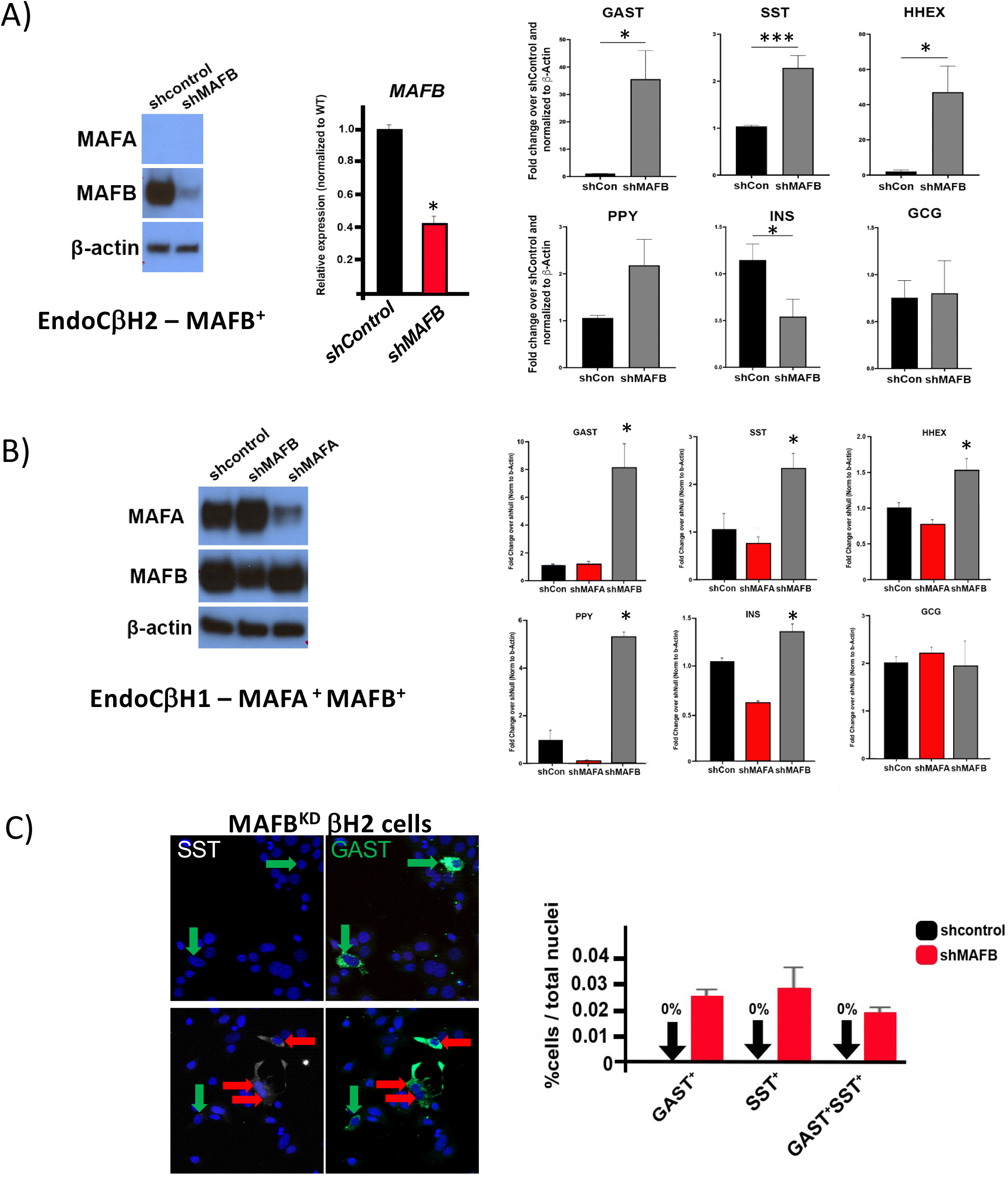
MAFB^KD^, but not MAFA^KD^, induces non-β cell hormone expression in human EndoC-β cells. Analysis of MAFA/B protein (left panels) and hormone-related mRNA levels (right panels) in shControl (scrambled construct), shMAFA- or shMAFB-treated (A) EndoC-βH2 (i.e., only MAFB^+^) and (B) EndoC-βH1 (i.e., MAFA^+^/B^+^) cells. β-Actin served as the internal control. n=3-4 replicates. Mean ± SEM. *p<0.05, ***p<0.005. Statistical significance was determined by comparing the shMAFA/B-treated groups to the shControl-treated groups. (C) Immunostaining for SST (white) and GAST (green) in MAFB^KD^ EndoC-βH2 cells. These proteins were undetectable in shControl-treated cells. Left, green arrows denote GAST^+^ SST^−^ cells, while red arrows denote GAST^+^ SST^+^ cells. Blue, DAPI staining of nuclei. Right, quantification in MAFB^KD^ EndoC-βH2 cells compared to shControl, with the fraction of hormone^+^ cells shown relative to the total cell number. Mean ± SEM. n=4 replicates.

In contrast to MAFB, EndoC-βH1 cells deficient in MAFA did not generate a significant change in *GAST* or *SST* mRNA levels (**Figure 3B, right, red bars**), even though MAFA protein levels were decreased more effectively than MAFB (**Figure 3B, left**). However, the reduction in MAFA did decrease *INS* and *PYY* mRNA production in MAFA^KD^ EndoC-βH1 cells. Collectively, our data suggests that β cell MAF TFs prevent non-β cell hormone gene expression in a species-specific manner, with only MafA serving in this capacity in mice and MAFB principally in humans.

### MAFB^KD^ EndoC-βH2 cells produce a gene signature reminiscent of the GAST^+^ MAFB^KO^ β-like cells

Sixty-one up-regulated genes were found by single cell sequencing of the human GAST^+^ MAFB^KO^ β-like cells derived from hESC cells (**Figure 4A, Supp Table 2**). By GO pathway analysis, many of these genes were linked to insulin (or hormone) secretion and glucose homeostasis (**Figure 4B**). Roughly 33% of the mouse S961-induced Gast^+^ cell signature overlapped with this human gene signature (7 of 20 genes, **Figure 4C**), while 34% of this dataset overlapped with genes enriched in *bona fide* GAST^+^ human stomach G cells (21 of 109 genes, **Figure 4D**) (28). However, only 5% of the gene changes were shared between these three populations (i.e., 3 of 61; CHGA, CHGB, and TTR; **Figure 4C-D**). Consequently, there are common and unique gene signatures found upon comparing the Gast^+^ cells generated pathologically from mouse β cells and human MAFB^KO^ GAST^+^ cells, and normal human stomach G cells.

**Figure 4:**
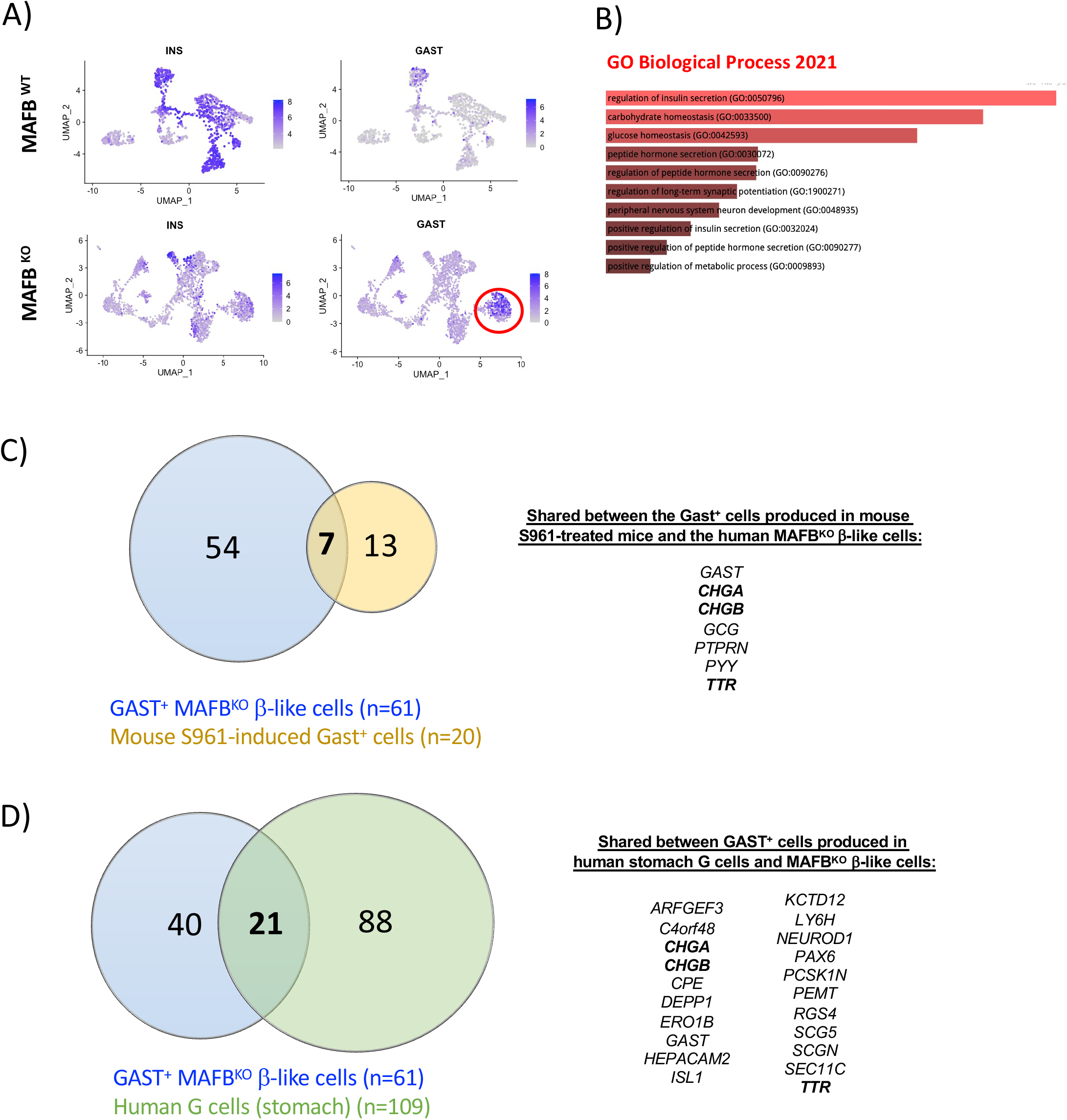
Unique and common gene signatures are produced in the Gast^+^ cells of mouse S961-treated β cells, hES-derived MAFB^KO^ β-like cells, and *bona fide* stomach G cells. (A) UMAP of the single-cell RNA-seq data from hESC-derived MAFB^WT^ and MAFB^KO^ cells ^7^. The number of GAST^+^ cells (encircled) was increased and INS^+^ cell numbers compromised in the MAFB^KO^ cell population relative to MAFB^WT^ cells. (B) Top 10 most influenced GO biological processes based on the 61 up-regulated genes in GAST^+^ MAFB^KO^ cells compared to GAST^−^ MAFB^KO^ cells. Venn diagrams of DEGs comparing single cell data from GAST^+^ cells from hESC-derived MAFB^KO^ β-like cells and (C) Gast^+^ β cells of mouse S961-treated islets, as well as (D) human stomach G cells. Gene products common to both conditions are listed on the right.

RNA sequencing of MAFB^KD^ EndoC-βH2 cells was performed next for comparison to the single cell datasets from hES-derived MAFB^KO^ GAST^+^ β-like cells, mouse S961-induced islet Gast^+^ cells, and human stomach G cells. Approximately 30% (18 of 61) of the up-regulated genes in MAFB^KO^ GAST^+^ β-like cells were enriched within the broader MAFB^KD^ EndoC-βH2 cell gene set (765 total up-regulated genes), several of which were validated by qPCR (**Figure 5A-C**). However, just 14% (15 of 109) of MAFB^KD^ EndoC-βH2 enriched genes overlapped with those found in human G cells (**Figure 5D**), with simply *DEPP1, HEPACAM2, PCSK1N*, and *RGS4* present in all three human GAST^+^ cell contexts (**Figure 5D**). Interestingly, non-hormone genes normally expressed in other human islet endocrine cell types, such as TMOD1 (γ and PP cells), PEG10 (α, γ, and PP cells), PCSK1N (α and PP cells), and HEPACAM2 (δ, γ, PP cells) (29), were produced in both human MAFB^KO^ GAST^+^ β-like cells and MAFB^KD^ EndoC-βH2 cells. In fact, PCSK1N, HEPACAM2, and RGS4 are also expressed in human stomach G cells (**Figure 5D**). Remarkably, *GAST* was the only common signature gene between the datasets from mouse S961-derived Gast^+^ cells, MAFB^KD^ EndoC-βH2 cells, hES MAFB^KO^ GAST^+^ β-like cells, and human G cells (**Supplemental Figure 3**). Together, our data not only demonstrates significant molecular diversity between normal Gast^+^ G cells and this hormone producing cell population generated under and pathophysiological conditions in the islet, but also highlights the importance of mouse MafA and human MAFB in regulating these species-specific gene signatures within the broader islet β cell population.

**Figure 5:**
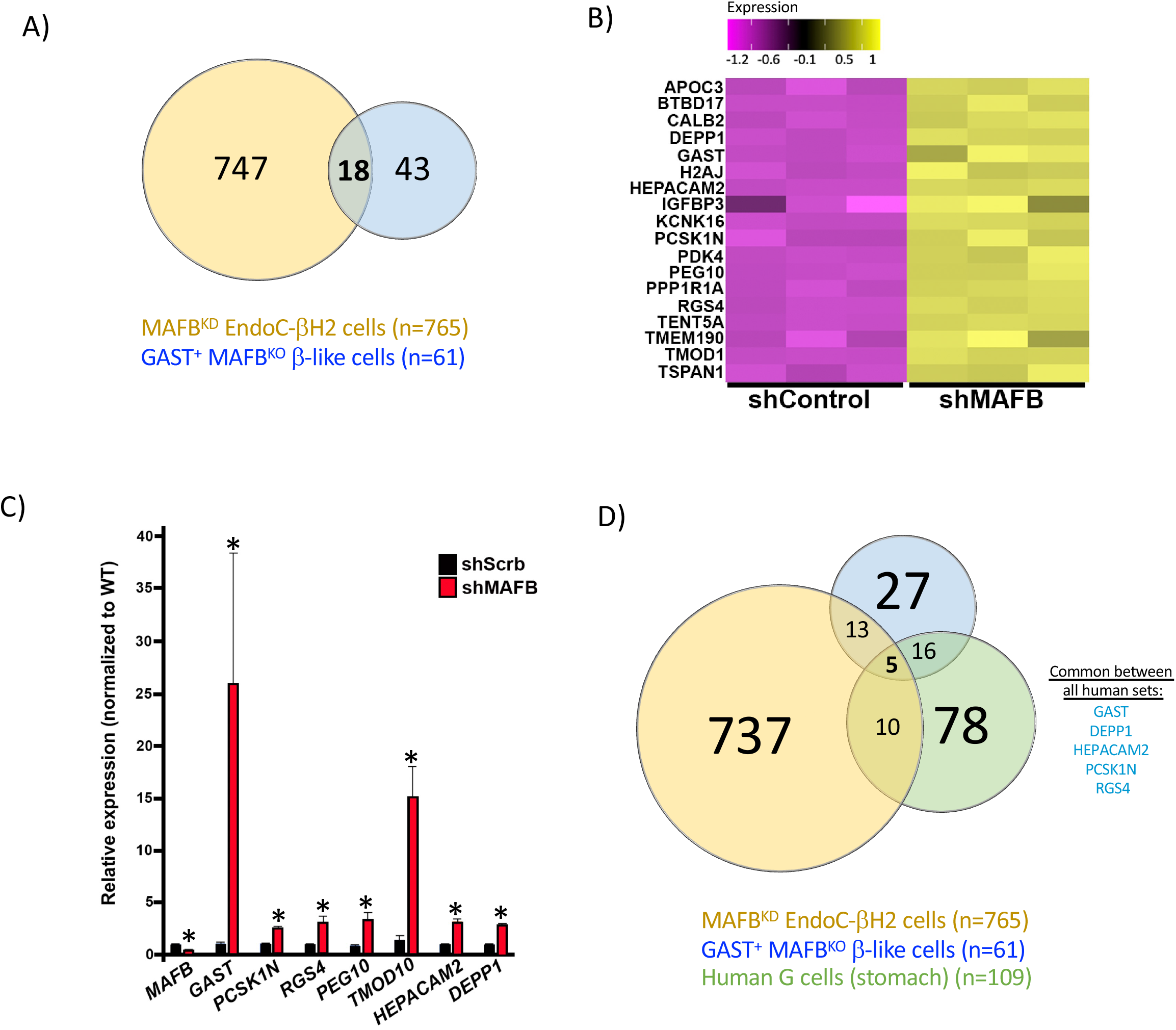
GAST^+^ MAFB^KO^ cell genes upregulated in MAFB^KD^ EndoC-βH2 cells. (A) Venn diagram of DEGs showing that 18 up-regulated genes are shared between MAFB^KD^ EndoC-βH2 cells and the GAST^+^ MAFB^KO^ hESC-derived β-like cells. (B) Heatmap of these 18 up-regulated shared genes in shMAFB- and shControl-treated EndoC-βH2 cells. (C) qPCR analysis of a subset of the shared genes in shMAFB- and shControl-treated EndoC-βH2 cells. Mean ± SEM. *p<0.05. n=3 replicates. (D) Venn diagram of DEGs of GAST^+^ MAFB^KO^ β-like cells, MAFB^KD^ EndoC-βH2 cells, and human stomach G cells. The 5 genes common to all three datasets are listed on the right

### T2D GAST^+^ cells are deficient in MAFB

Both MAFA and MAFB are highly sensitive to glucotoxicity and oxidative stress, with protein levels reduced in T2D islets (30), (31). To determine if MAFB levels were low in the GAST^+^ cells produced in T2D islets (27), serial sections from normal and T2D donors were immunostained for both proteins. Indeed, GAST was only detectable in T2D tissues in relatively low MAFB-producing (MAFB^LOW^) cells (**Figure 6A-C, Supplemental Table 3**). GAST^+^ cells appeared to also be produced in female T2D samples (**Supplemental Figure 4**). Additionally, rare GAST^+^SST^+^ dual positive cells were observed in the survey of male T2D samples, a pattern found in EndoC-βH2 cells (**Figure 3D**) and previously in T2D donors (27).

**Figure 6:**
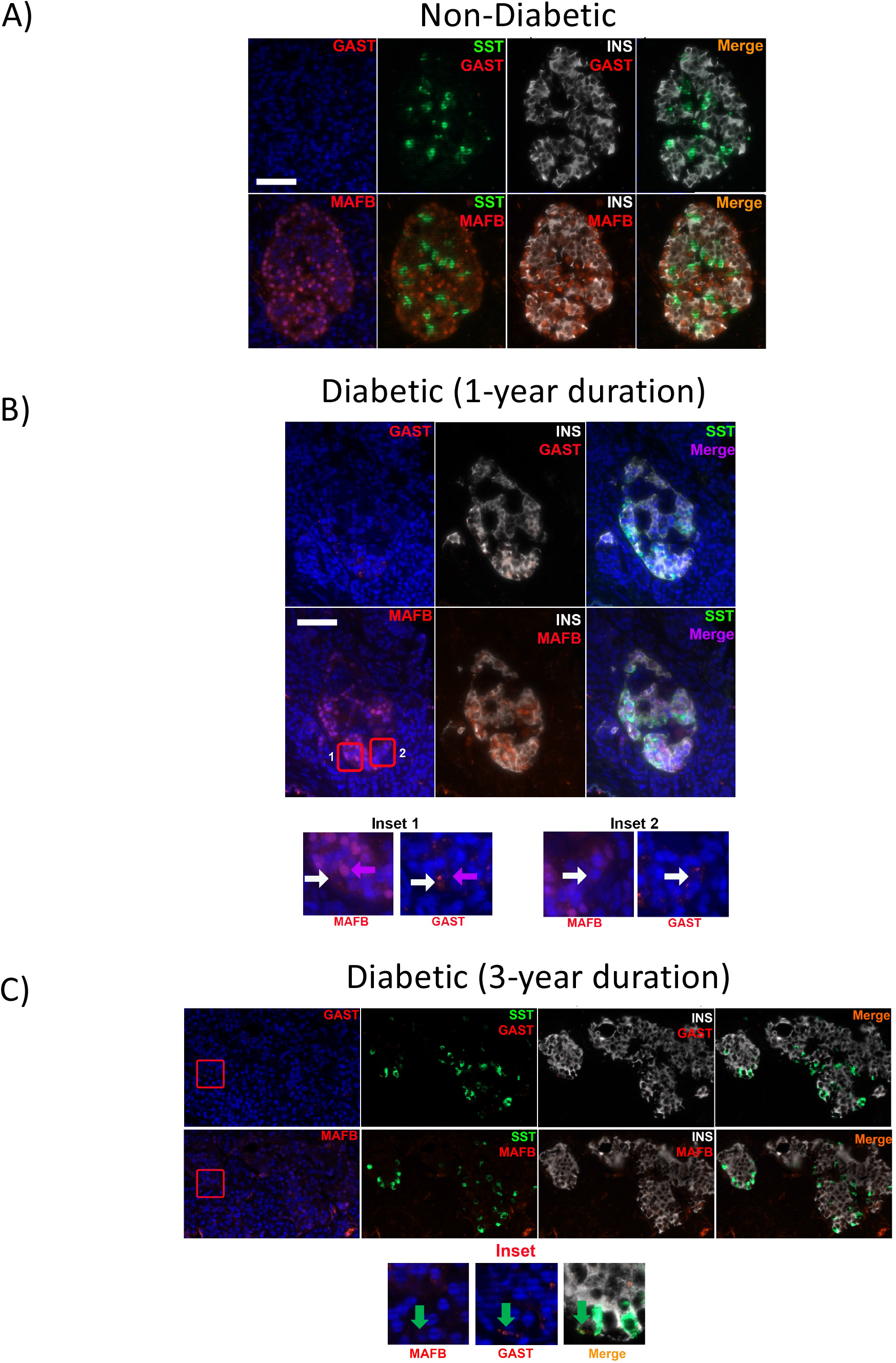
GAST is produced in MAFB^LOW^ T2D islet cells. (A-C) Representative images of immunostaining performed on serial sections from male non-diabetic and diabetic pancreata: GAST (Red in top panels), MAFB (Red in bottom panels), SST (Green), INS (White), and nuclei (blue). (A) GAST^+^ cells were not detected in healthy donor islets by immunofluorescence analysis. However, GAST production was associated with lower MAFB levels in T2D islet cells from donors at (B) 1- and (C) 3-years from diagnosis. Bar, 100 µm. Insets show magnified view of representative GAST^−^ MAFB^HI^ (Purple arrows), rare GAST^+^ MAFB^LOW^ (White arrows), and GAST^+^ MAFB^LOW^ SST^+^ (Green arrows) cells.

To further analyze the relationship between human *GAST* gene expression and MAFB, *GAST* mRNA levels were measured in human islets following either U6- or rat *Insulin* enhancer/promoter (i.e., RIP)-driven lentiviral-mediated *MAFB* knockdown (pseudoislets). *GAST* was elevated after a roughly 50% *MAFB* knockdown in these contexts (**Supplemental Figure 5**). These results further support a role for MAFB in repressing *GAST* expression in adult human islet β cells.

### MAFB directly regulates human *GAST* transcription

Because human islet MAFB binding sites were detected approximately 1.5 kb upstream of the *GAST* gene transcriptional start site by ChIP-Seq (32, 33), we asked whether this TF could directly bind and control β cell gene transcription (**Figure 7A**). Notably, the presence of other islet-enriched TFs (i.e., PDX1, NKX2.2, FOXA2, and NKX6.1) in this region indicted these may also act in concert with MAFB to negatively or positively impact expression (32).

**Figure 7:**
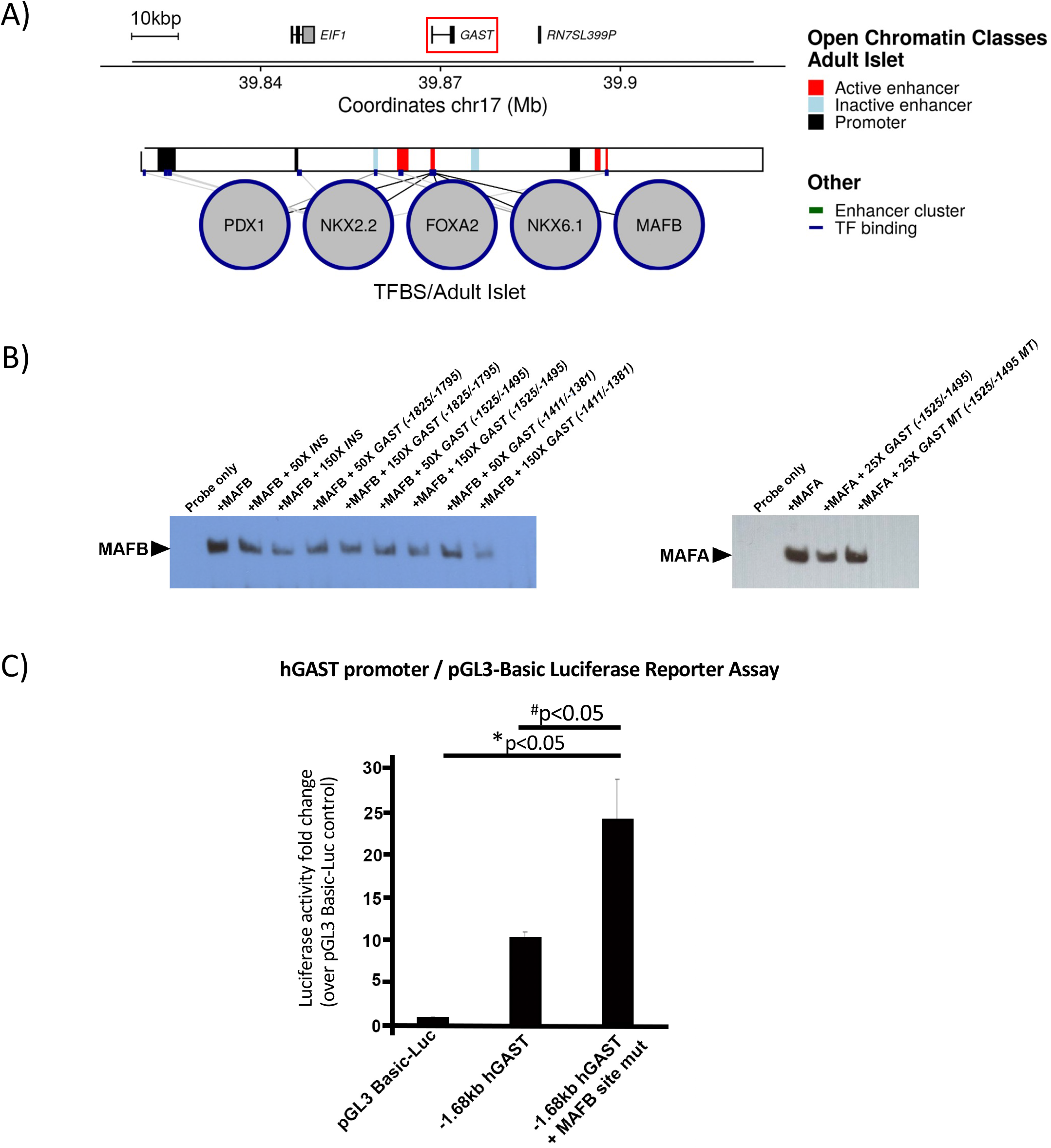
MAFB binds within the human *GAST* 5’-flanking region and suppresses gene expression in β cells. (A) ChIP-seq data derived from whole islets by Pasquali et al ^32^ illustrating islet-enriched PDX1, NKX2.2, FOXA2, NKX6.1 and MAFB TF binding sites within approximately 1.5 Kbp of the human islet *GAST* gene transcription start site. (B) Gel shift analysis of MAFB (left) or MAFA (right) protein binding to a biotin-labeled human *INSULIN (INS)* probe to a MAFA/B binding site starting at position -135 in the presence of human *INS* and human *GAST* -1825/-1795, -1525/-1495, or -1411/-1381 bp unlabeled competitors. While all three *GAST* sites competed as effectively as the *INS* element, only the *GAST* -1525/1495 element is conserved in mouse and also binds MAFA. n=3-4 replicates. (C) MAFA/B binding site mutation in the -1525/-1495 element stimulated -1.68 kbp *GAST*-driven luciferase reporter activity in human EndoC-βH1 cells. pGL3 Basic-LUC represents the control vector without insert. n=3 replicates. Mean ± SEM. *p<0.05 compared to Basic-Luc control; ^#^p<0.05 compared to hGAST control.

The ability of endogenous MAFB to bind to 5’-flanking sequences in the human *GAST* gene was analyzed in gel shift assays with EndoC-βH1 nuclear extracts. This region appears to contain three MAFB binding sites (i.e., -1825/-1795, -1525/-1495, and -1411/-1381), as indicated by their ability to compete effectively in relation to a well characterized MAFA/MAFB binding site in the human *INS* gene (**Figure 7B, left**). Only one of these human MAFB binding sequences is conserved in mouse (i.e., -1525/-1495), which also effectively bound to MAFA (**Figure 7B, right**). As predicted for a repressor binding site, mutation of this conserved element prevented the inhibitory effect of MAFB and led to elevated expression of the -1.68 kb-driven *GAST* reporter in transfected EndoC-βH2 cells (**Figure 7C**).

## DISCUSSION

MAFB was not only shown to be essential for insulin production during the formation of hESC-derived β-like cells but also in preventing non-β cell hormone expression (7). Here we analyzed if similar regulatory properties were found for MafA during the formation of mouse islet β cells, and MAFA and/or MAFB in adult human β cells. To this end, we compared the gene regulatory properties of these TFs across multiple conditions, including direct genetic manipulation of MafA in mouse islet β cells, MAFA^KD^ and MAFB^KD^ in human EndoC-βH1/2 β cells, MAFB^KO^ in hES-derived β-like cells, and MAFB^KD^ in human pseudoislets as well as in pathophysiologic conditions in mouse models and human T2D islets. Our experimental results revealed that: (1) the genes activated by insulin resistance in Gast^+^ cells are also expressed in the mouse *MafA*^*Δβ*^ and *ob/ob* islet β cell population, and not in normal islet α, δ, or PP cells, (2) MAFB^KD^, but not MAFA^KD^, induces GAST^+^ cell production in EndoC-βH1/2 cells, demonstrating species-specific regulation by these TFs, (3) the GAST^+^ cells produced in MAFB^KO^ hESC-derived β-like cells or pathologically in mouse models are distinct from physiologic human stomach G cells, (4) readily detectible Gast protein production is restricted to a small subpopulation of MafA/MAFB-deficient β cells, however the genes enriched in these cells are represented in the broader TF-depleted β cell populations, and (5) MafA and MAFB appear to directly repress *GAST* transcription in β cells. As the levels of MAFA and MAFB are more sensitive to metabolic stressors in relation of other islet-enriched TFs (30), their compromised expression upon exposure to obesity- and insulin resistance-induced effectors could lead to β cell dysfunction early and preceding development of T2D.

Of the many peptide hormones expressed by the mouse pancreatic islet, gastrin is uniquely restricted to the fetal pancreas. Pancreatic Gast^+^ cells are derived from Ngn3 TF^+^ endocrine progenitors, are abundant during development but wane at birth, and are undetectable in adult pancreata (27). Deletion of *Gast* from the mouse pancreas does not affect islet cell composition or the islet proliferative response to injury (34). However, gastrin has been proposed to represent an endocrine cell reprogramming marker, with production representing a reversible cell state (20). Consequently, GAST^+^ cells may represent a pool of “reprogrammable” cells of loosened identity which can be coaxed back to β cells or produce new β cells from non-β cell sources in an autocrine and/or paracrine manner. Several groups have attempted to harness its properties to expand the β cell compartment (35-40). If this is the case, it is prudent to understand how similar GAST^+^ cells produced from human MAFB^KO^ hES β-like cells or pathophysiologically in the setting of MAFB deficiency are to *bona fide*, physiologic human GAST-expressing cells. Unfortunately, the molecular composition of embryonic GAST^+^ cells of the pancreas is unknown, although there is ∼33% identity between the GAST^+^ cells generated from human MAFB^KO^ hES β-like cells to human stomach G cells, and to mouse S961-treated β cells (**Figure 4C,D**). However, very few of these genes are shared between the mouse and human GAST^+^ cell populations (i.e., 5%, **Figure 4C,D**). Hence, while the GAST^+^ cells produced by insulin resistance in mice or the MAFB^KO^ GAST^+^ β-like cells have some molecular similarities to stomach G cells, these conditions alone are insufficient to completely reprogram the β cell towards a *de facto* G cell.

Strikingly, while Gast protein was only detected in a small number of cells in *MafA*^*Δβ*^ mouse islets (**Figure 1C**), S961-treated mouse islets (**Figure 2C**), and MAFB^KD^ EndoC-βH2 cells (**Figure 3C**), many of the genes detected were found throughout the bulk RNA-seq analyzed β cell populations of *MafA*^*Δβ*^ islets (**Figure 2D**), *ob/ob* islets (**Supplemental Figure 2A**) and MAFB^KD^ EndoC-βH2 cells (**Figure 5B**). Even though the genes comprising the S961-induced Gast^+^ signature include other islet cell hormones (**Supplemental Table 1**), *MafA*^*Δβ*^ β cell gene expression was more reminiscent of this Gast^+^ cell population than prototypical islet α, δ, and PP cells (**Figure 2E**). This data indicates that β cell dysfunction under insulin resistant and/or obese circumstances results from induction of a Gast^+^/GAST^+^ cell-like molecular phenotype due to loss of MafA in mice and MAFB in humans.

The heterogeneity amongst Gast^+^/GAST^+^ cells produced in mouse *MafA*^*Δβ*^ islets, human MAFB^KD^ EndoC-βH1/2 cells, and human G cells could reflect a variety of factors. For example, baseline β cell heterogeneity in MafA/MAFB and calcium/calcineurin signaling may contribute, as deficiency in both are involved in GAST^+^ β cell production (27). In addition, this heterogeneity could entail experimental context (i.e., animal versus cell line) and/or differences in temporal induction of Gast^+^ cell associated genes in relation to loss to TF expression.

Because MAFA and MAFB recruit the mixed-lineage leukemia 3 (MLL3) and MLL4 coregulator complex to alter chromatin structure and gene expression (5), we analyzed whether *GAST* expression was regulated by this effector in MAFB^KD^ EndoC-βH2 cells. However, knockdown of a core subunit of these methyltransferases, NCOA6, did not influence MAFB^KD^-regulated *GAST* expression (**Supplemental Figure 6**). Future studies using established methods to identify the coregulator(s) mediating the repressive effects of MafA and MAFB in β cells (5). Importantly, is possible that continual repression of non-β cell transcriptional programs are another function of islet-enriched TFs that are presently associated with gene activation. For example, deficiency in other key islet TFs permit misexpression of non-β cell hormones, such as Gcg, Sst, Ppy, Pyy, and Ghrelin. These results reveal the significance of such TFs in controlling intra-islet cell plasticity (reviewed in (11)). However, Gast^+^ cells were not produced upon down regulation of the Pdx1(41), Nkx6.1(42), Nkx2.2(43), or Pax6(44), consistent with possibility of a unique pattern of cell identity control by such TFs (11). It is noteworthy that MafA mutant mice have a subtler islet phenotype than these other islet-enriched TF mutants (45), thus, correlating with earlier inducers of β cell dysfunction such as obesity.

More robust *Gast* expression was identified in male mouse islets than female in hyperglycemic mice and *MafA*^*Δβ*^ mice, suggesting a greater vulnerability of male β cells to metabolic stressors and alternative cell states. This is supported by the higher rates of overt diabetes in males than females in mouse models of diabetes and multiple forms of human diabetes (reviewed in (46)). In addition, transplantation of hESC-derived β-like cells into female recipient mice often yield faster in vivo maturation and insulin secretion properties than male recipient mice (47). Interactions between the MafA, MafB, c-Maf and nuclear sex hormone receptors (i.e., estrogen receptor β and androgen receptor) warrant further study, especially given the results described here and sex-dependent effects produced by the S64F MAFA variant (48). While our studies demonstrate GAST^+^ cells in both male (**Figure 6**) and female (**Supplemental Figure 4**) T2D donors, comprehensive studies using more donors are needed to discern possible sex differences in the regulation of human islet *GAST*, GAST-related peptides (such as CCK (**Figure 1A**)), and stress induced *GAST* cell-associated gene products.

Although the presence of GAST^+^ cells in human T2D islets was recognized earlier (27), the relevance to diabetes pathogenesis was unclear. Our results strongly suggest that GAST^+^ gene signatures are enriched in metabolically stressed, but pre-diabetic, islet β cells. Moreover, the enrichment of these signature genes throughout an islet population with a very limited number of overt GAST^+^ cells further implicate an increased sensitivity to MafA or MAFB protein levels. As a result, we predict that these associated gene signatures will be induced even earlier than GAST itself upon pathologically induced loss of MafA or MAFB, which could be tested temporally in future studies in models of insulin resistance (i.e., following S961- or HFD-treatment). It is also unclear how these differentially expressed genes influence islet β cell dysfunction or identity under these conditions. Additionally, we recognize that several of the islet GAST^+^ cell enriched signature genes are normally expressed in other islet endocrine cells, thus, determining if differences in their regulation occur will require detailed investigation across normal, prediabetic, and diabetic settings using β cell enriched experimental methods. In fact, GAST^+^ δ cells have been detected in human T2D, although MAFB is not typically expressed in this cell type. Here, we find that SST is induced in a MAFB-deficient, pure β cell line (**Figure 3C**); however, we cannot exclude the possibility of δ cell reprogramming in an in vivo setting. Along this line, MAFB is robustly expressed in islet α cells, which raises the question how the compromised levels of this TF in pathogenic settings (e.g., T2D (30)) impacts these cells.

In summary, this work further highlights alternative β cell states imposed by metabolic stressors. This work puts forth a novel regulatory role of islet MafA and MAFB in maintaining mono-hormonal β cell identity by actively repressing non-β Gast^+^ cell-associated transcriptional programs. Early loss of MAFB in T2D may promote β cell reprogramming, and understanding the stepwise transitions towards dysfunction, which are potentially reversible, may facilitate the production of therapies in which early β cell dysfunction can be targeted to slow the progression to overt diabetes. Future studies to understand the natural history and production of Gast^+^ and other non β cell hormone^+^ cells (e.g., Cck), namely by lineage tracing under physiologic and pathophysiologic conditions and identifying strategies to target these cells will be critical in our understanding of alternative β cell fates and potentially coaxing their return to normal functional β cells.

## ACKNOWLEDGEMENTS

This research was performed using resources and/or funding provided by NIH grants to R.S. (DK090570), J.C. (K08 DK132507), E.M.W. (F32 DK109577), M.Huising (R01 DK110276), M.Hebrok (R01 DK090570) and by the Vanderbilt Diabetes Research and Training Center (DK20593). J.C. by a Doris Duke Charitable Foundation Physician Scientist Fellowship (2020063) and Burroughs Wellcome Fund Career Award for Medical Scientists; X.T. was supported by a JDRF Fellowship (3-PDF-2019-738-A-N) and Diabetes Research Connection (2022); A.M. by the Stephen F. and Bettina A. Sims Immunology Fellowship and the AWS Machine Learning Research Award; M.H. by a JDRF award (2-SRA-2021-1054-M-N); and Y.D. by DKG. The human pancreatic sections were generously provided by Alvin Powers and Marcela Brissova from the Vanderbilt Pancreas Biorepository that was created with the support of DK106755, DK108120, DK104211, and the Vanderbilt Diabetes Research and Training Center (DK020593). We also thank Drs. Raphaël Scharfmann and Phillippe Ravassard for providing EndoC-βH1/2 cells, Drs. Alberto Delgado and Keith Wilson for providing mouse stomach tissues, and Dr. Daniel Turkewicz for technical support. S961 was provided as a gift from Novo Nordisk to the Dor lab. Imaging was performed with NIH support from the Vanderbilt University Medical Center Cell Imaging Shared Resource (NCI grant CA68485; NIDDK grants DK20593, DK58404, and DK59637; NICHD grant HD15052; and NEI grant EY08126).

## Author Contributions

JC, XT, EW, RS designed the study.

JC, XT, EW, TD, VC, SA, RR, AO, AM, MG, JL performed experiments.

JC, XT, EW, TD, VC, SA, RR, AO, AM, performed bioinformatic analysis.

JC, XT, EW, TD, VC, SA, RR, AO, AM, MG, JL, MH, MM, MH, YD, RS analyzed data.

JC, XT, RS wrote the manuscript.

JC, XT are co-first authors due to their contributions in most of the experiments described. JC was listed first for having a larger role in writing the manuscript.

## METHODS AND MATERIALS

### *MafA*^*Δβ*^ generation, S961 treatment, and blood glucose measurements

*MafA*^*Δβ*^ mice were generated by crossing floxed *MafA* (*MafA*^*fl/fl*^)(3) with *Ins1*^CreERT2^ (*Ins1* ^tm1(CreERT2)Thor^) knock-in mice(15). *MafA*^*fl/f*^;*Ins1* ^CreERT2^ (*Ins1* ^tm1(CreERT2)Thor^) mice were then crossed with *Ins2*^*Apple*^ mice (49). For isolation of β-cells, islets from Apple producing mice were sorted by FACS on the red spectra to an average purity of 85–95% using the Vanderbilt Flow Cytometry Shared Resource core. The S961-treated insulin resistant mouse model was established as previously described (27). Briefly, vehicle (PBS) or 12 nmol S961 (Insulin receptor antagonist, a gift from Novo Nordisk to the Dor lab) was loaded into an ALZET osmotic pump (model 2001) and implanted subcutaneously on the back of 6-week-old male ICR (CD-1) mice for 4-7 days. Random blood serum glucose measurements were collected through the tail vein without fasting.

### Single-cell RNA sequencing and analysis of S961-treated mouse islets

Control and S961-treated diabetic mouse islets were isolated, dissociated and stained with propidium iodide (PI). Live cells (PI negative) were sorted using FACS Aria III (BD Biosciences). Libraries were made using 10X Genomics Chromium Single Cell 3ʹ Reagent Kit V3 following manufacturer instructions and sequenced on Illumina HiSeq 2500. Raw reads of each sample were processed using the ‘count’ command of the Cell Ranger software, v2.0.2, aligning the reads to the mouse mm10 (GRCm38) genome version. The generated report was used for assessing the quality of the samples in terms of cell numbers (3072, 2443), average reads per cell (104108, 149852), fraction of reads in cells (92.3%, 93.6%), alignment rate and saturation (57.9%, 61.3%) for PBS- and S961-treated groups, respectively. The cells of the PBS (i.e., 3072)- and S961 (2443)-treated groups were further filtered to leave only high-quality cells according to the criteria of having between 500 to 4500 genes, less than 50000 UMIs and less than 5% of mitochondrial RNA. Genes expressed in less than 3 cells were removed. 2984 and 2355 cells, for PBS- and S961-treated cells, respectively, were retained for a further analysis using the package Seurat 3.0.2 on R3.6.3, as previously described (50).

### Islet isolation, shRNA-based knockdown, RNA isolation, cDNA synthesis, and real-time PCR

Mouse islets were isolated using collagenase P (Roche Applied Science) injected into the pancreatic duct, followed by a Histopaque (1119 and 1077; Sigma-Aldrich) gradient. Islets were incubated overnight in standard RPMI-1640 medium (Biological Industries) supplemented with 10% FBS, L-glutamine, and penicillin-streptomycin in a 5% CO_2_ incubator at 37°C. Hand-picked islets were subject to RNA isolation and real-time qPCR analysis. The scrambled shRNA (VectorBuilder, Chicago, IL), MAFA shRNA (Genecopoeia, Rockville, MD), and MAFB shRNA (VectorBuilder, Chicago, IL) lentiviruses were grown in human embryonic kidney 293T cells and concentrated using the PEG-it Virus Precipitation Solution (System Biosciences, Mountain View, CA), as previously described (51). The knockdown experiments were performed in EndoC-βH1/2 cells (25, 26). Cells were infected for 36h and followed by 1:5000 puromycin selection for 36-42h. RNA was then isolated using the RNeasy micro kit (Qiagen). Mouse islet RNA was collected using the TRIzol reagent (Life Technologies) and a DNA-free RNA kit (Zymo Research). cDNA was generated using Superscript III reverse transcriptase (Invitrogen) by the oligo(dT) priming method. Real-time PCR assays were performed using the LightCycler FastStart DNA Master PLUS SYBR Green kit (Roche) and a LightCycler PCR instrument (Roche). The real-time PCR primers are listed in **Supplemental Table 4**. Real-time PCR results were analyzed using the ΔΔCt method; *Gapdh* or *β-Actin* was used to normalize the data.

### Human pseudoislets

Human islets (**Supplemental Table 3**) were obtained from the IIDP (https://iidp.coh.org/), Alberta Diabetes Institute Islet Core (https://www.epicore.ualberta.ca/IsletCore/), and the Human Pancreas Analysis Program (https://hpap.pmacs.upenn.edu/). Primary human islets were cultured in CMRL 1066 (MediaTech, 15-110-CV media (5.5 mM glucose), 10% FBS [MilliporeSigma, 12306C], 1% Penicillin/Streptomycin [Gibco, 15140-122], 2 mM L-glutamine (Gibco, 25030-081]) in 5% CO_2_ at 37°C for ∼24 hours before beginning the studies. Briefly, human islets were handpicked to >95% purity and then dispersed with 0.025% HyClone trypsin (Thermo Scientific). Islet cells were counted and transfected with shRNA lentiviral vectors at an MOI of 500. Infected cells were seeded at 2000 cells per well in Cell-Carrier Spheroid Ultra-low attachment microplates (PerkinElmer) in enriched Vanderbilt pseudoislet media (52). Cells were allowed to reaggregate for 6 days before being harvested for RNA isolation and qPCR analysis. Donor information is listed in **Supplemental Table 3**.

### EndoC-βH1 and EndoC-βH2 cells

Human EndoC-βH1 and EndoC-βH2 cells were maintained under conditions described previously (25, 26). The growth medium contains DMEM (Gibco and Thermo Fisher Scientific), 5.6 mmol/L glucose, 2% BSA (Serologicals Proteins), 100 mU/mL penicillin, 100 mg/mL streptomycin, 50 mmol/L 2-mercaptoethanol, 10 mmol/L nicotinamide, 5 mg/mL transferrin, and 6.7 ng/mL sodium selenite (SigmaAldrich, St. Louis, MO). EndoC-βH1/2 cells were infected for 1-2 weeks with 150 ng shControl, shMAFA, or shMAFB lentiviral particles/million cells before harvesting for immunoblotting and qPCR analysis. Knockdown for human NCOA6 was performed by siRNA transfection (5) (Dharmacon) for 48h before harvesting for qPCR analysis. MAFA and MAFB protein levels were normalized to endogenous β-actin by immunoblotting with anti-MAFA (Cell Signaling 79737), anti-MAFB (Cell Signaling 41019S) and anti-β-actin (4967S, Cell Signaling) antibodies. Horseradish peroxidase-conjugated anti-rabbit or anti-goat secondary antibodies were used at 1:5000. Immunoblots were quantitated with ImageJ software (National Institutes of Health). Antibody information can be found in **Supplemental Table 5**. The *GAST-*driven firefly luciferase constructs were transfected in EndoC-βH1 cells along with phRL-TK internal control (Promega) using the Lipofectamine protocol (Life Technologies). *GAST-*driven firefly luciferase spans sequences from -1680 bp to +8 bp, and the -1680 bp to +8 bp and -1525/-1495 element mutant were constructed in the pGL3-Basic luciferase reporter vector (Promega) using standard molecular biology techniques. Cellular extracts were collected 48h post-transfection, and the Dual-Luciferase Reporter Assay (Promega) performed according to the manufacturer’s directions. Firefly luciferase measurements were normalized to the Renilla phRL-TK internal control.

### Electrophoretic mobility shift assay

HeLa cells were transfected with either a CMV-driven MAFA or MAFB expression plasmid using the Lipofectamine protocol. Forty-eight hours later, nuclear extract and DNA binding reactions were performed. Briefly, 10μg of nuclear extract and 200fmol of the biotin-labeled double strand human *Insulin* MAFA/B binding site probe were mixed either alone or with unlabeled competitor DNAs in a 20μl reaction system (LightShift Chemiluminescent EMSA Kit, Thermo Scientific) containing 1X binding buffer, 2.5% glycerol, 5 mM MgCl, 50 ng/ul of poly(dI-dC), and 0.05% NP-40. MAFA and MAFB antibody super-shift reactions were performed to localize the TF:probe band. The sequences of the probe and competitors is provided in **Supplemental Table 6**. The reactions were separated on a 6% pre-cast DNA retardation gel in 0.5% Tris borate-EDTA buffer (TBE) at 100 V for 1.5 h.

### Immunofluorescence staining

Human pancreata, mouse pancreata and treated EndoC-βH1/2 cells were fixed in 4% paraformaldehyde in PBS on ice (6h fixation for pancreata, 15min fixation on slides for cells), and pancreata were embedded in either Tissue-Plus O.C.T. (Thermo Fisher Scientific) or paraffin wax. The cryosections of human and rodent pancreas as well as fixed EndoC-βH1/2 cells were made permeable by 0.5% Triton treatment for 10 min. The paraffin sections were de-paraffinized and rehydrated before citrate buffer-based antigen retrieval. Following blocking with 0.5% BSA in PBS for 120 min, the primary antibodies were applied overnight at 4°C. Species-matched antibodies conjugated with the Cy2, Cy3, or Cy5 fluorophores were used for secondary detection (1:1,000; Jackson ImmunoResearch, West Grove, PA). Immunofluorescent images were obtained using the Zeiss Axio Imager M2 widefield microscope with ApoTome. Primary antibodies are listed in **Supplemental Table 5**. DAPI was used for nuclear staining (Southern Biotech).

### Bulk RNA-Sequencing and analysis

The RNeasy micro plus kit (Qiagen) was used to isolate total RNA from FACS-purified mouse islet β cells and EndoC-βH2 cells (n≥3 independent replicates). Isolated RNA quality was analyzed on an Agilent 2100 Bioanalyzer. Only samples with an RNA Integrity Number >8.0 were used for library preparation. cDNA libraries were constructed and paired-end sequencing performed on an Illumina NovaSeq6000 (150-nucleotide reads). The generated FASTQ files were processed and interpreted using the Genialis visual informatics platform (https://www.genialis.com) as described before (48). Gene expression levels were quantified with HTSeq-count, and differential gene expression analyses performed with DESeq2. Poorly expressed genes, which have expression count summed over all samples below 10, were filtered out from the differential expression analysis input matrix. RNA expression analysis of selected candidates was performed with the qPCR primers provided in **Supplemental Table 4**.

### Statistics

Significance was evaluated with the Student two-tailed t-test or one-way ANOVA, with statistical value denoted in the figure legends.

### Study approval

Mice of both sexes were used in this study; details can be found in the results section and figure legends. All animal studies were reviewed and approved by the Vanderbilt University Institutional Animal Care and Use Committee. Mice were housed and cared for according to the Vanderbilt Department of Animal Care and the Institutional Animal Care and Use Committee of Animal Welfare Assurance Standards and Guidelines. Human pancreata from healthy and T2D donors were received for analysis in partnership with the International Institute for the Advancement of Medicine National Disease Research Interchange Integrated Islet Distribution Program and nPOD. Donor information is listed in **Supplemental Table 3**. The Vanderbilt University institutional review board declared that studies on deidentified human pancreatic specimens do not qualify as human subject research.

## FIGURE LEGENDS

**Supplemental Figure 1: Effective *Mafa* deletion in *MafA*^*Δβ*^ mice**

(A) Immunostaining for MafA (red) and Ins (white) in 3-month-old male WT and *MafA*^*Δβ*^ pancreata. n=3 animals per group.

(B) *Mafa* mRNA levels in WT and *MafA*^*Δβ*^ mouse islets. n=3-4 animals per group. Mean ± SEM. *p<0.05.

**Supplemental Figure 2. The S961-induced Gast^+^ gene signature is maintained in male, but not female, *ob/ob* mouse islets**.

(A) The presence of elevated Gast^+^ signature genes were determined by analysis of the RNA-seq datasets prepared from FACS-sorted β cells of lean female (n=5 animals), lean male (n=5 animals), *ob/ob* female mice (n=2 animals), and *ob/ob* male mice (n=2 animals). The islet Gast^+^ genes from S961-treated mice (**Supp. Table 1**) largely overlapped with those in male and not female *ob/ob* mouse islet β cells.

(B) Heatmap of Gast^+^ signature genes in female *MafA*^*Δβ*^ islets. n=4 animals/group.

**Supplemental Figure 3: Venn diagram analysis of the DEGs expressed in Gast^+^ cells of S961-treated mouse islets, human MAFB^KO^ β-like cells, human stomach G cells with the broader MAFB^KD^ EndoC-βH2 cell population**.

Only Gast itself was enriched in all four datasets.

**Supplemental Figure 4: GAST is produced in female MAFB^LOW^ T2D islet cells**.

(A-B) Representative image of immunostaining performed on serial sections to detect GAST (Red in top panels), MAFB (Red in bottom panels), SST (Green), INS (White), and nuclei (blue). GAST^+^ cells were not detected in female healthy donor islets but was in MAFB^LOW^ T2D islets. Bars, 50 µm. Insets show magnified view of rare GAST^+^MAFB^**LOW**^ (White arrows) and GAST^−^MAFB^**HI**^ (Purple arrows) cells.

**Supplemental Figure 5: *GAST* is increased in MAFB^KD^ human pseudoislets**.

(A-B) qPCR analysis of whole pseudoislets (n=2 donors) showed increased *GAST* expression upon targeting MAFB either in all islet cells (shMAFB-U6) or only β cells (shMAFB-RIP) in relation to the scramble control (shControl).

**Supplementary Figure 6: MLL3/4 does not influence MAFB-mediated repression of *GAST* expression in EndoC-βH2 cells**.

Representative qPCR analysis of EndoC-βH2 cells subject to MAFB and/or NCOA6 knockdown by shRNA and siRNA, respectively. Reduction of the core MLL3/4 subunit, NCOA6, did not accentuate *GAST* upregulation by MAFB reduction. n=5 replicates. Mean ± SEM. *p<0.05.

**Supplemental Table 1: Mouse Gast^+^ β cell signature (male) table**

**Supplemental Table 2: Human GAST^+^ β cell signature (male) table**

**Supplemental Table 3: Human islet donor characteristics table**

**Supplemental Table 4: qPCR primer table**

**Supplemental Table 5: Primary antibody table**

**Supplemental Table 6: Competitor probe sequences table**

