## Supplemental Figures for "Species-specific roles for the MAFA and MAFB transcription factors in regulating islet β cell identity"

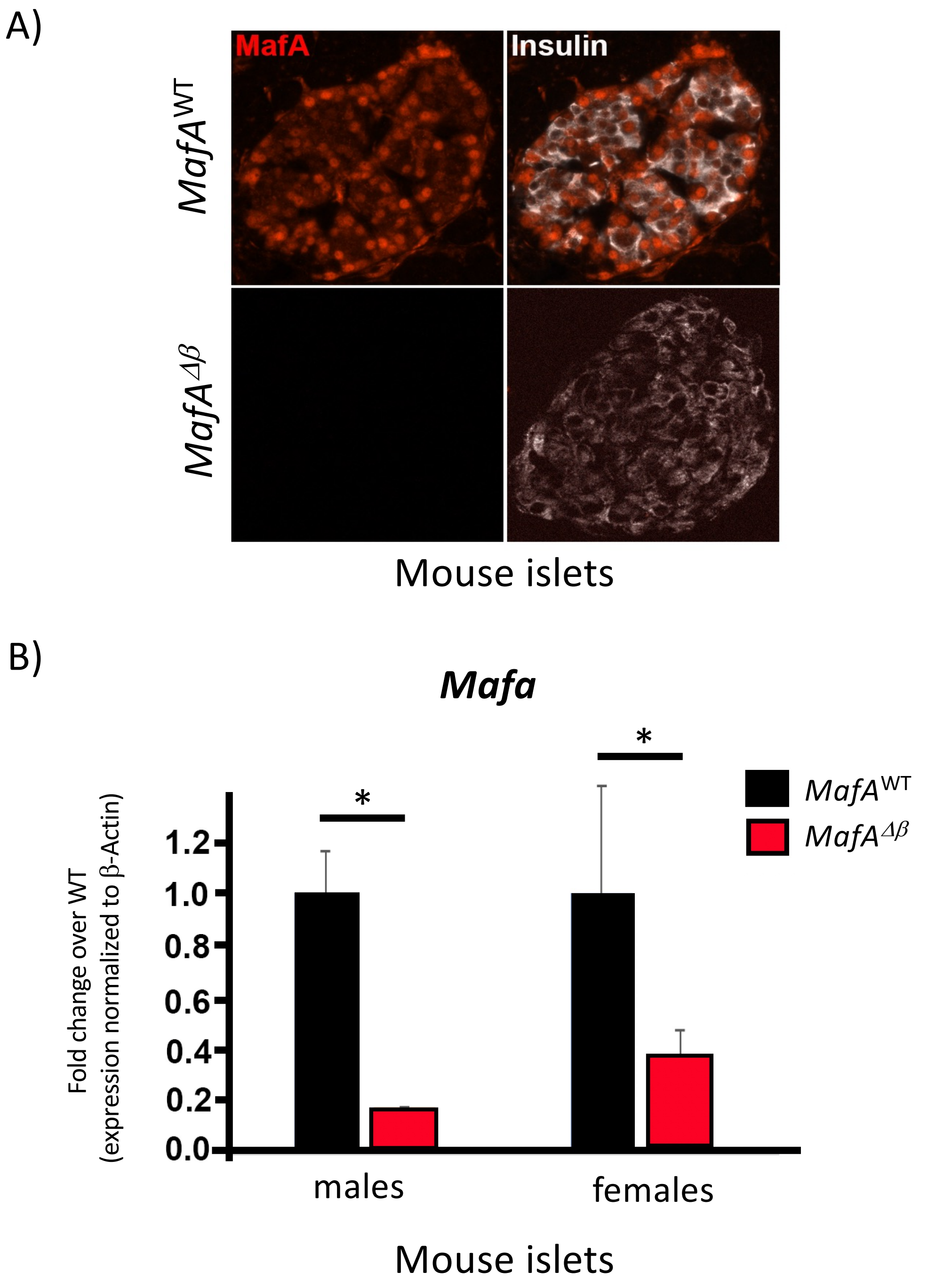


**Supplemental Figure 1: Effective *Mafa* deletion in *MafA*$^{\Delta\beta}$ mice**
(A) Immunostaining for MafA (red) and Ins (white) in 3-month-old male WT and *MafA*$^{\Delta\beta}$ pancreata. n=3 animals per group.
(B) *Mafa* mRNA levels in WT and *MafA*$^{\Delta\beta}$ mouse islets. n=3-4 animals per group. Mean ± SEM. *p<0.05.

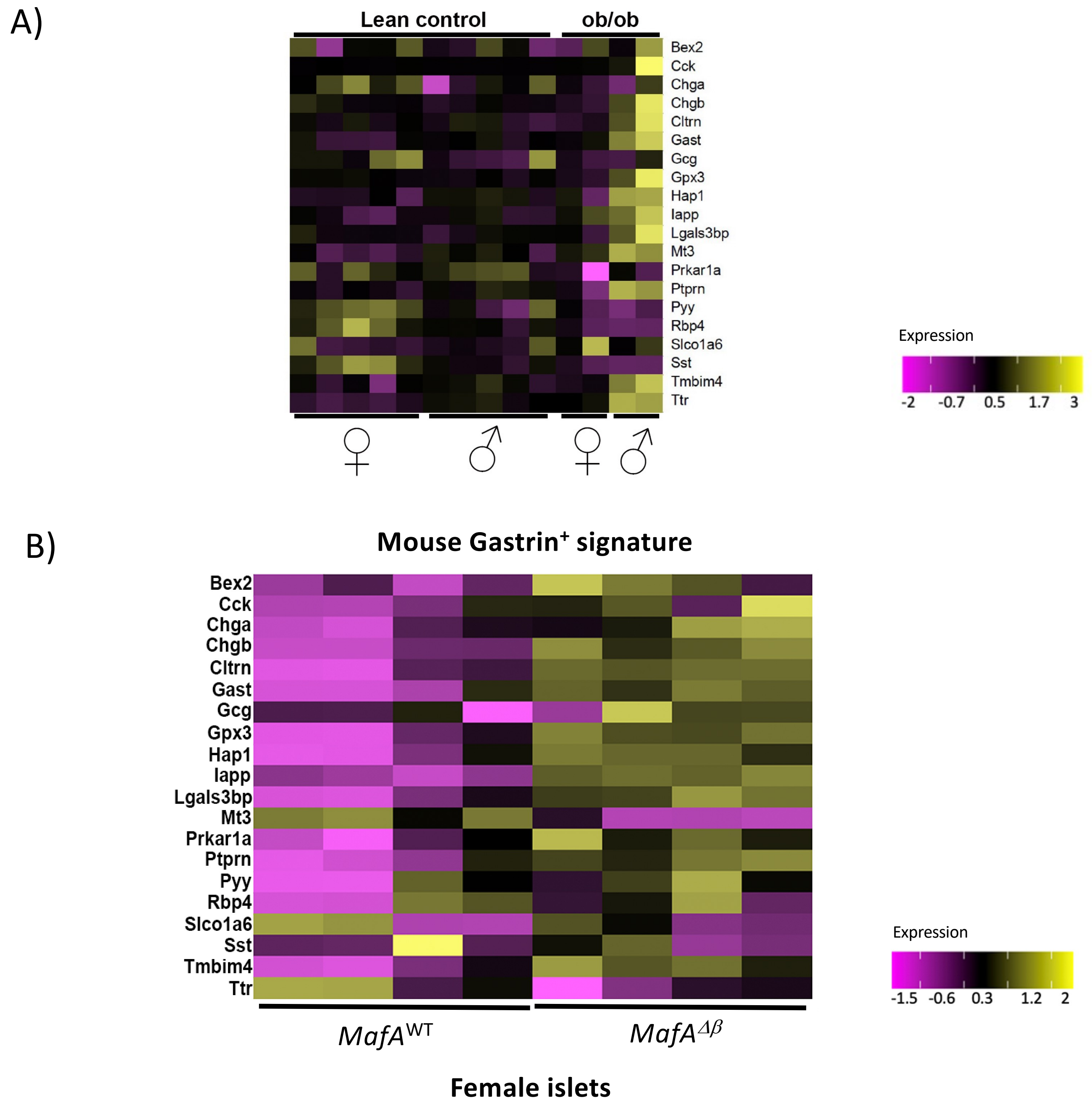


**Supplemental Figure 2. The S961-induced Gast+ gene signature is maintained in male, but not female, *ob/ob* mouse islets.**

(A) The presence of elevated Gast+ signature genes were determined by analysis of the RNA-seq datasets prepared from FACS-sorted β cells of lean female (n=5 animals), lean male (n=5 animals), *ob/ob* female mice (n=2 animals), and *ob/ob* male mice (n=2 animals). The islet Gast+ genes from S961-treated mice (**Supplemental Table 1**) largely overlapped with those in male and not female *ob/ob* mouse islet β cells.

(B) Heatmap of Gast+ signature genes in female *MafA*$^{\Delta\beta}$ islets. n=4 animals/group.

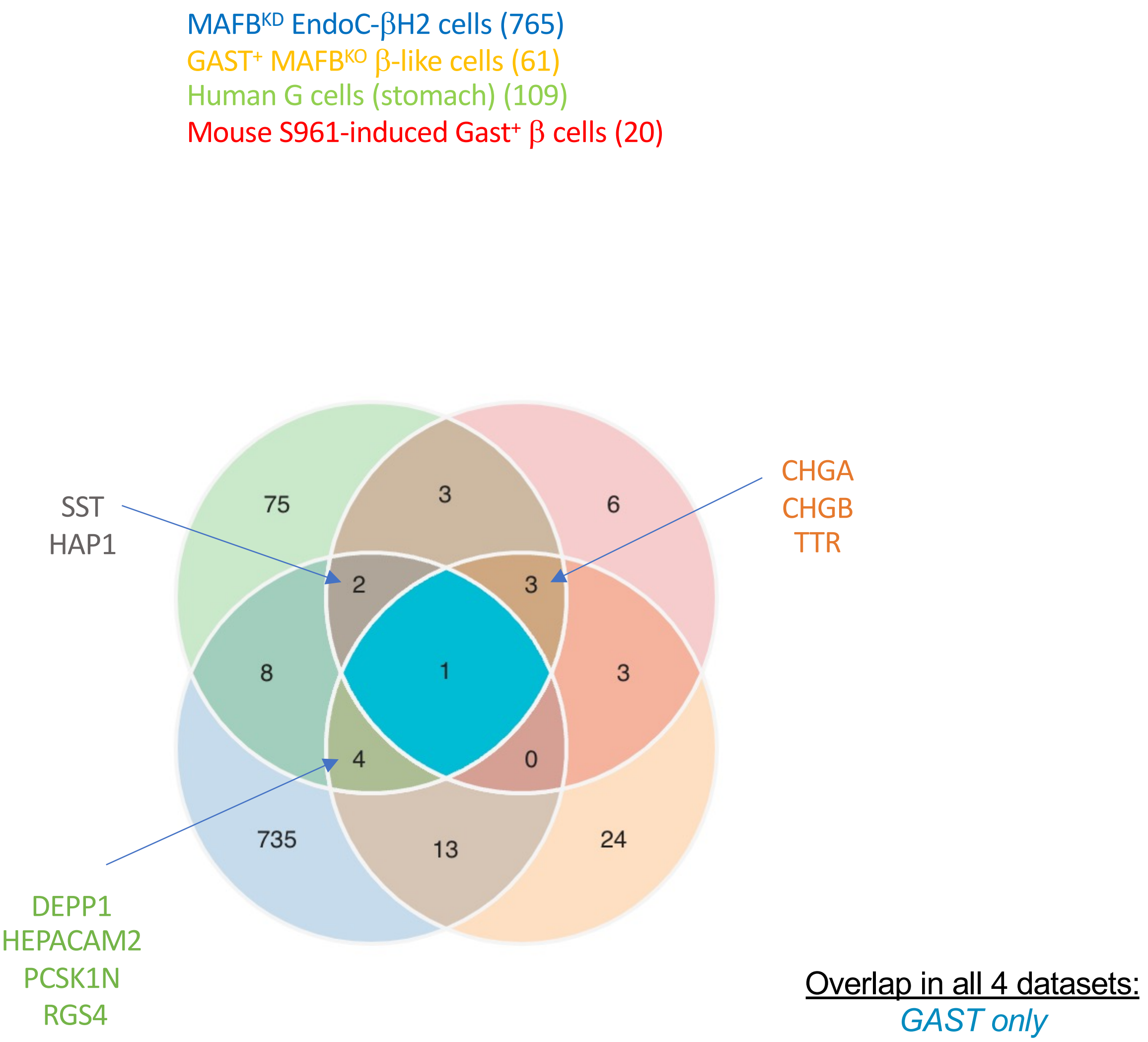


**Supplemental Figure 3: Venn diagram analysis of the DEGs expressed in Gast$^{+}$ cells of S961-treated mouse islets, human MAFB$^{KO}$ β-like cells, human stomach G cells with the broader MAFB$^{KD}$ EndoC-βH2 cell population.** Only Gast itself was enriched in all four datasets.

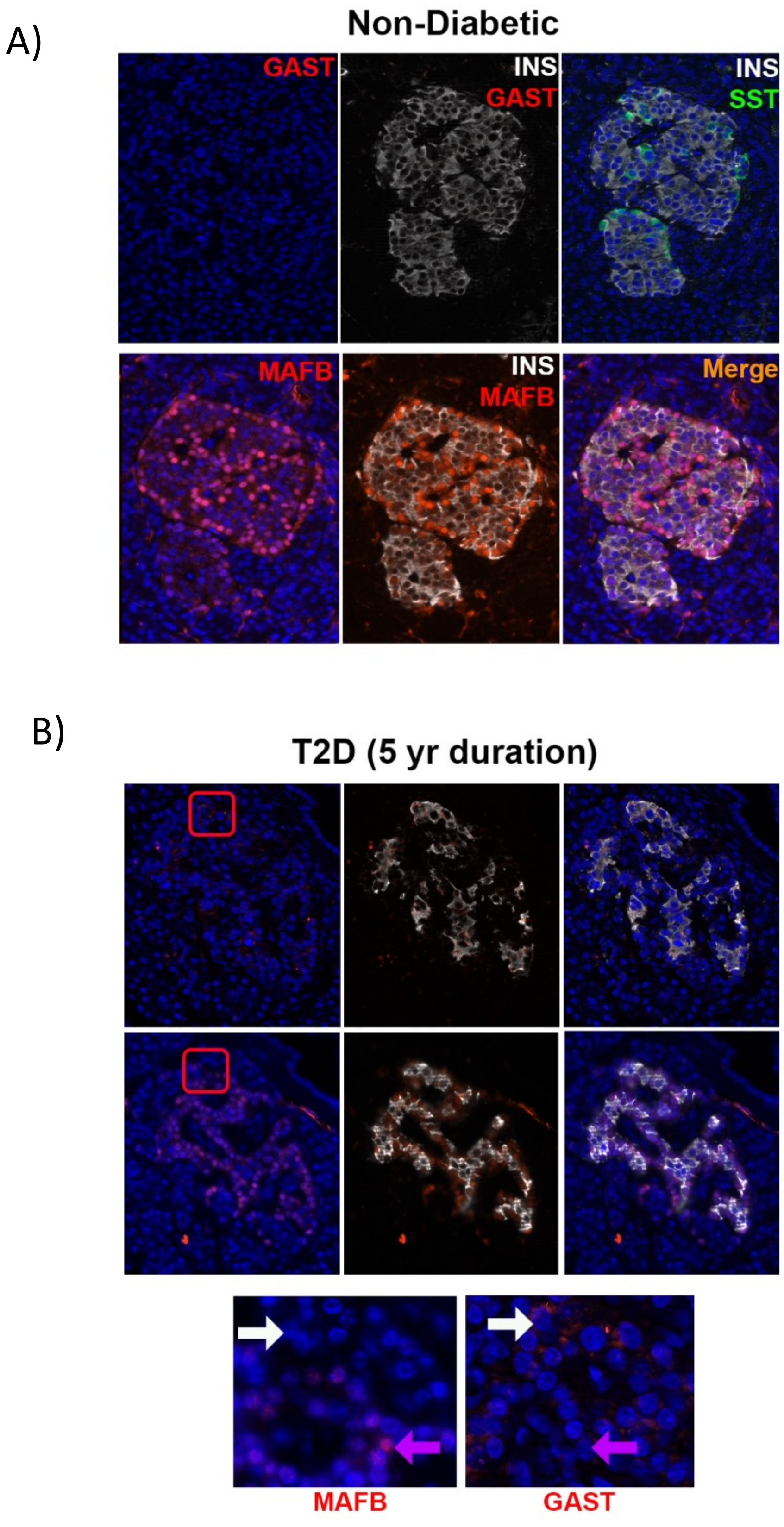


**Supplemental Figure 4: GAST is produced in female MAFB$^{LOW}$ T2D islet cells.**

(A-B) Representative image of immunostaining performed on serial sections to detect GAST (Red in top panels), MAFB (Red in bottom panels), SST (Green), INS (White), and nuclei (blue). GAST$^{+}$ cells were not detected in female healthy donor islets but was in MAFB$^{LOW}$ T2D islets. Bars, 50 μm. Insets show magnified view of rare GAST$^{+}$MAFB$^{LOW}$ (White arrows) and GAST$^{-}$MAFB$^{HI}$ (Purple arrows) cells

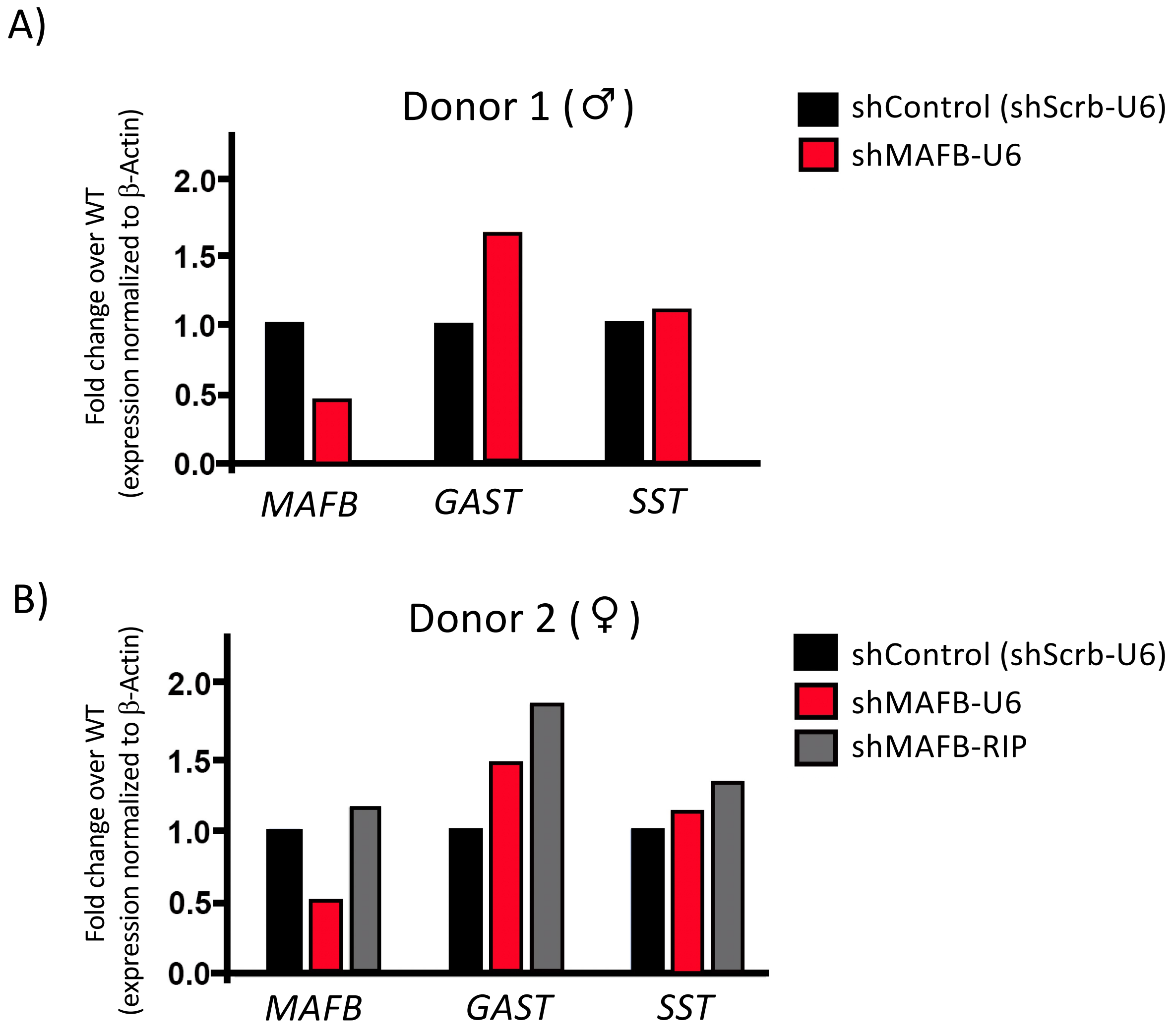


**Supplemental Figure 5: *GAST* is increased in MAFB$^{KD}$ human pseudoislets.**
(A-B) qPCR analysis of whole pseudoislets (n=2 donors) showed increased *GAST* expression upon targeting MAFB either in all islet cells (shMAFB-U6) or only β cells (shMAFB-RIP) in relation to the scramble control (shControl).

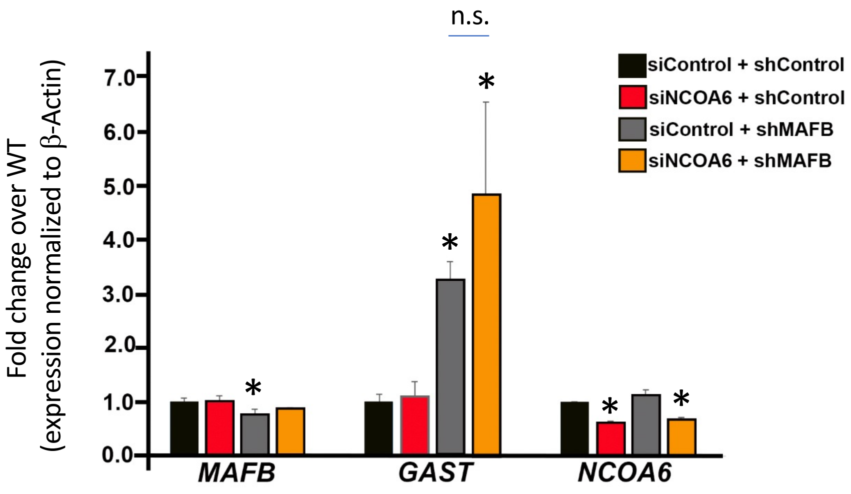


**Supplementary Figure 6: MLL3/4 does not influence MAFB-mediated repression of *GAST* expression in EndoC-βH2 cells.**

Representative qPCR analysis of EndoC-βH2 cells subject to MAFB and/or NCOA6 knockdown by shRNA and siRNA, respectively. Reduction of the core MLL3/4 subunit, NCOA6, did not accentuate *GAST* upregulation by MAFB reduction. n=5 replicates. Mean ± SEM. *$p<0.05$.
